# Challenges and opportunities in population monitoring of cheetahs

**DOI:** 10.1101/563122

**Authors:** Daniel W. Linden, David S. Green, Elena V. Chelysheva, Salim Mandela, Stephanie M. Dloniak

## Abstract

Population monitoring is key to wildlife conservation and management but is challenging at the spatial and temporal extents necessary for understanding changes. Non-invasive survey methods and spatial capture-recapture (SCR) models have revolutionized wildlife monitoring by providing the means to more easily acquire data at large scales and the framework to generate spatially-explicit predictions, respectively. Despite the opportunities for improved monitoring, challenges can remain in the study design and model fitting phases of an SCR approach. Here, we used a search-encounter design with multi-session SCR models to collect spatially-indexed photographs and estimate the changes in density of cheetahs between 2005 and 2013–2016 in the Masai Mara National Reserve (MMNR) in southwestern Kenya. Our SCR models of cheetah encounters suggested little change in cheetah density from 2005 to 2013–2016, though there was some evidence that density fluctuated annually in the MMNR. The sampling period length (5 vs. 10 months) and timing (early, late, full year) over which spatial encounters were included in the modeling did not substantially alter inferences about density when sample sizes were adequate (>20 spatially distinct encounters). We estimated an average cheetah density of ~1.2 cheetahs/100 km^2^, consistent with the impression that the MMNR provides important habitat for cheetahs in Africa. During most years and seasonal periods, the spatial distribution of vegetation greenness (a proxy for ungulate habitat quality) accounted for important variation in encounter rates. The search-encounter design used here could be applied to other regions for the purposes of cheetah monitoring. While snap-shot estimates of population size across time are useful for wildlife monitoring, open population models could identify the mechanisms behind changes and further facilitate better conservation and management decision making.

## Introduction

Population monitoring is key to wildlife conservation and management but is challenging at the spatial and temporal extents necessary for understanding changes (Ellis et al. 2014). Monitoring over space and time requires a feasible scheme and persistence in both dedication and resources to obtain adequate information. Low replication in either dimension reduces the capacity to explain observed patterns or test hypotheses about perturbation, limiting the value of the monitoring data for informing conservation and management decisions (Yoccoz et al. 2001). The monitoring challenge has been particularly acute for wide-ranging, cryptic species that occur at low densities, such as carnivores. These life history features have historically made data collection and analysis difficult and reduced the opportunities for robust inference about population dynamics at relevant spatial and temporal scales (Karanth et al. 2006).

Non-invasive survey methods (Long et al. 2008) and spatial capture-recapture (SCR) models (Borchers and Efford 2008, Royle et al. 2014) have revolutionized wildlife population monitoring by providing the means to more easily acquire data at large scales and the framework to generate spatially-explicit predictions, respectively. In an SCR model, the locations of individual encounters (e.g., photographs, genetic material) are used to determine centers of activity for each observed individual, providing spatial information on the number of total individuals in the population and the probabilities of encountering them across the landscape. By formally linking the distributions of individuals and their movement ecology in a hierarchical framework, SCR models jointly estimate the ecological and observational processes that generate the spatial encounter data collected by large-scale monitoring designs, enabling robust inferences that are critical for conservation (Royle et al. 2018). These models have proven useful for estimating the density of wide-ranging carnivores, particularly in applications to large felids including tigers *Panthera tigris* (Royle et al. 2009), jaguars *Panthera onca* (Sollmann et al. 2011), leopards *Panthera pardus* (Gray and Prum 2012) and cougars *Puma concolor* (Russell et al. 2012). Recently, the approach was illustrated using search-encounter surveys with African lions *Panthera leo* (Elliot and Gopalaswamy 2017) and cheetahs *Acinonyx jubatus* (Broekhuis and Gopalaswamy 2016). These applications have highlighted the potential of SCR as a monitoring tool, though rarely have studies spanned long enough timeframes to allow for examining temporal changes in population size or density at large scales (e.g., Chandler and Clark 2014).

Despite the opportunities for improved monitoring, challenges can remain in the study design and model fitting phases of a spatial capture-recapture approach. Sampling efforts may not yield enough unique spatial locations per individual to enable model fitting (Becker et al. 2017), unless some type of auxiliary data is integrated (e.g., telemetry; Sollmann et al. 2013). Longer survey durations can be used to acquire more captures or encounters, at the expense of potentially violating assumptions regarding population closure (i.e., no births, deaths, immigrants/emigrants during sampling). The timing and duration of surveys will dictate the scope of the population being assessed, dependent on which individuals are available for sampling (e.g., residents vs. dispersers) and can meet assumptions of the observation process. Resource selection at one or more spatial scales can affect model inferences if not properly incorporated, particularly if it results in unmodeled heterogeneity in the encounter process (Royle et al. 2013, Linden et al. 2018). And small sample sizes, even when large enough to enable model fitting, may yet afford little power for accommodating relevant variation in one or more parameters which can reduce accuracy and precision of the resulting estimates (Sollmann et al. 2013). Most of these design and modeling considerations are important for any animal sampling and population estimation approach, and we note that explicitly modeling the sampling process does not necessarily obviate critical assumptions regarding how data were collected and what the data represent. For these reasons and others, it is prudent that researchers design robust monitoring schemes, use multiple lines of evidence, and temper any conclusions from monitoring data when making inferences that will guide conservation and management of large carnivore populations.

Here, we used a search-encounter design with SCR models (sensu Royle et al. 2011) to collect spatially-indexed photographs and estimate the changes in density of cheetahs between 2005 and 2013–2016 in the Masai Mara National Reserve (MMNR) in southwestern Kenya. Cheetahs are currently listed globally as “vulnerable” with a decreasing total population (Durant et al. 2015, Durant et al. 2017) and while much of the current cheetah range exists outside of protected areas the populations within represent important strongholds for cheetah conservation (Durant et al. 2017). Few long-term studies have empirically estimated how cheetah populations are faring over time (Chauvenet et al. 2011, Durant et al. 2011), or have illustrated how changing landscapes around protected areas may be influencing wildlife within reserve boundaries. Carnivore populations in the MMNR have historically been high compared to other areas in sub-Saharan Africa (Craft et al. 2015), and the Mara-Serengeti ecosystem is considered a stronghold for large carnivores in East Africa (Ogutu and Dublin 2002, Riggio et al. 2013). Yet, populations of wild herbivores in the MMNR have been declining over time (Ottichilo et al. 2000, Ogutu et al. 2009, Ogutu et al. 2011), livestock often graze within reserve boundaries and anthropogenic disturbance has altered the behaviors and population numbers of other large carnivores (Boydston et al. 2003, Kolowski and Holekamp 2009, Green et al. 2018a), and rangelands around the MMNR are rapidly shifting into a matrix of urbanization and agriculture (Lamprey and Reid 2004, Løvschal et al. 2017).

Broekhuis and Gopalaswamy (2016) recently provided a 2014 population estimate for cheetahs within the greater Mara using a similar survey and SCR modeling approach. We fit more extensive data from a multi-year survey effort (2005, 2013–2016) conducted during a longer sampling window (10 months) with sample sizes that afforded additional model complexity. In particular, we incorporated a resource selection function relating the probability of encounter to annual variance in green vegetation (i.e., Normalized Difference Vegetation Index [NDVI]) as an approximation to habitat quality for ungulate prey (Pettorelli et al. 2005, Bro-Jorgensen et al. 2008). We hypothesized that cheetahs would be encountered more frequently in areas with high variation where vegetation changed drastically across the year in response to moisture (e.g., short grass), compared to low variance regions with relatively constant conditions (e.g., riparian forest or bare ground). We also compared inferences between 5-month (both an early and late season) and 10-month sampling periods to explore tradeoffs in the acquisition of encounters while trying to meet population closure assumptions.

Our earlier initial modeling efforts suggested a >50% decline in cheetah density between 2005 and 2013 (Green et al. 2014), but the population estimate by Broekhuis and Gopalaswamy (2016) challenged that conclusion. Additional years of monitoring and subsequent modeling indicate that the Mara cheetah population may exhibit annual fluctuations due to movement between the MMNR, adjacent conservancies, and the Serengeti National Park, highlighting the importance of conservation and management efforts in those areas surrounding the reserve.

## Materials and Methods

### Study area and data collection

Our study took place in the 1510 km^2^ Masai Mara National Reserve in southwestern Kenya (Figure 1). The MMNR is predominantly comprised of open grassland interspersed with riparian areas, supporting a high density and diversity of resident herbivores, which are also joined seasonally by migrant populations of wildebeest *Connochaetes taurinus*, zebra *Equus quagga*, and Thomson’s gazelle *Eudorcas thomsonii* from the Serengeti National Park to the southwest and the Loita plains to the northeast (Bell 1971, Stelfox et al. 1986, Sinclair and Norton-Griffiths 1995). The MMNR is bounded by the border with Tanzania and the Serengeti National Park to the south, and is surrounded in all other directions by community conservancies, pastoralist communities, small towns, and agricultural lands (Figure 1). There are no fences or barriers encompassing the MMNR, and wildlife regularly move beyond its political borders.

**Figure 1.**
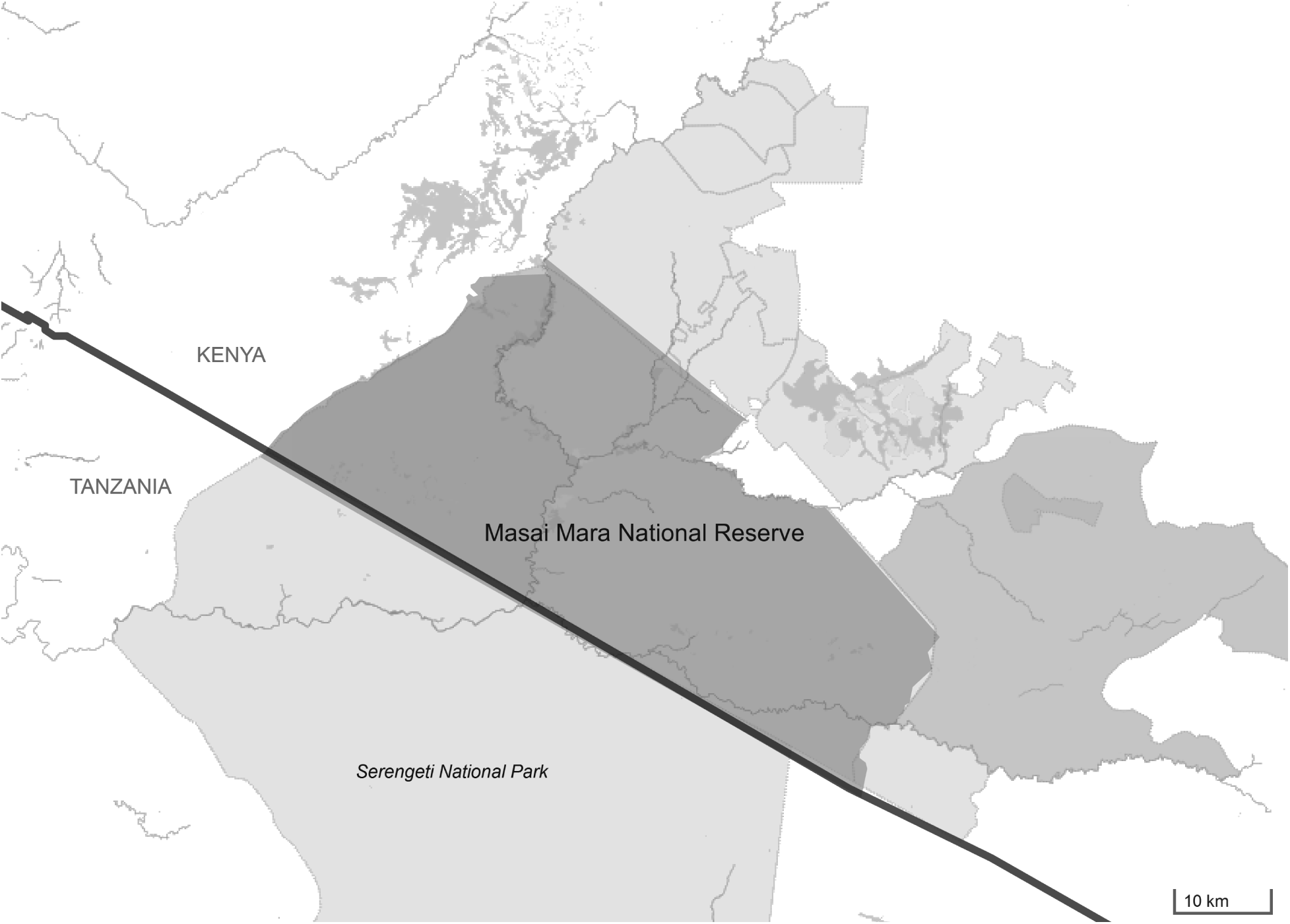
The location of cheetah monitoring in the Masai Mara National Reserve (MMNR) in southwestern Kenya (35.125° E, 1.44° S). Other conservation areas (shaded) surround the MMNR, including the Serengeti National Park in Tanzania to the south. Map data © OpenStreetMap contributors, CC BY-SA.

We systematically searched for cheetahs in the MMNR from January to October in 2005 and each year during 2013–2016 by dividing the MMNR into 6 sampling blocks roughly equal in size (Figure S1). Searches occurred between 0500 and 1900 h, during which time observers (1 or 2) drove throughout one block looking for cheetahs in a single vehicle, periodically stopping and scanning the surrounding landscape with binoculars (Caro 1994). Main roads were followed when convenient but considerable time was spent off-road to cover all accessible areas of each block; survey effort was calculated as the number of hours spent searching a block on a given date. When a cheetah was sighted, we drove within 50 m of an individual or group of individuals and photographed both sides of each animal and recorded geographic coordinates, sex and age class. We identified each individual using the distinct pelage and tail ring patterns (Caro and Durant 1991) and limited our modeling to adults.

We acquired spatial raster data from the Famine Early Warning System Network hosted by the USGS/EROS Data Center (https://earlywarning.usgs.gov/fews/). The data included 250 m resolution grids with 10-day NDVI values observed across each year (36 for a given year) for a region spanning most of East Africa. We calculated the standard deviation in NDVI value within a given year to approximate the seasonal variation within a given grid cell. Notable features that are apparent in every year include the vegetation along the Mara and Talek Rivers (Figure S2).

### Spatial capture-recapture model

Similar to previous applications of spatial capture-recapture using unstructured search-encounter designs (Russell et al. 2012, Broekhuis and Gopalaswamy 2016), we divided our study area (the MMNR) into a grid with a sufficiently low resolution (2-km × 2-km cells) to create spatial encounter histories for individual cheetahs. We defined the number of encounters *y_ij_* for individual *i* in grid cell *j* as a Poisson-distributed random variable:

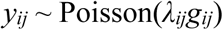

Here, *λ_ij_* is the mean encounter rate for an individual having its activity center (s_*i*_) within a given grid cell, and *g*_*ij*_ is a detection function describing how encounter rate decreases as the distance (*d*_*ij*_) increases between the location of an individual’s activity center and the coordinates of grid cell *j*. We chose a Gaussian encounter probability model such that *g_ij_* = exp(–*d*_*ij*_^2^/2*σ*^2^), where *σ* is a scale parameter representing the standard deviation of a bivariate normal distribution used to approximate space usage (Royle et al. 2014). While sex is often used as a factor for describing variation in *σ* (Sollmann et al. 2011, Broekhuis and Gopalaswamy 2016), our early model fitting did not indicate a difference between females and males or among years; *σ* remained constant in our final model specification.

The mean encounter rate *λ_ij_*was modeled as a function of several variables specific to an individual and grid cell. We considered differences among years to account for potential factors related to observers and the space-use of individual cheetahs in a given year. We also considered two grid cell covariates for *λ_ij_*: 1) the annual variance in NDVI for each year (standardized within the year to have mean = 0 and unit variance); and 2) the log-transformed fraction of hours spent searching a grid cell, given its location within 1 of the 6 search blocks. We included quadratic functions for NDVI that were year-specific to accommodate resource selection by cheetahs in response to spatial-temporal differences in vegetation within the Mara across years. The effect of search effort was constrained similar to a Poisson offset, though we estimated a regression coefficient instead of assuming it was 1. As such, we modeled the log-linear encounter rate (*λ_ij_*) as:

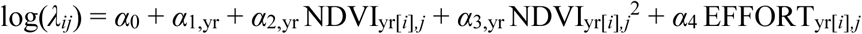

Here, *α*_0_ is the baseline encounter rate on the log scale for an individual captured in 2005; *α*_1,yr_ is a vector of year-specific coefficients for differences in encounter rates of individuals observed in later years (yr[*i*] = 2013, 2014, 2015, or 2016); *α*_2,yr_ and *α*_3,yr_ are vectors of year-specific coefficients for the linear and quadratic effects, respectively, of the variance in NDVI for each grid cell in each year; and *α*_4_ is a coefficient describing the relationship between encounter rate and search effort in a given grid cell and year. We considered encounters separated by ≥5 days to represent independent events with regards to individual movement and encounter probability and, therefore, thinned 18–35% of the total encounters in a given year to help meet model assumptions. Adult male cheetahs regularly form coalitions with other males (Caro and Collins 1987) and we observed them doing so in the MMNR (~60% of male sightings involved coalitions). Despite this, we treated each sighting as an independent observation given that coalitions were sometimes observed to exhibit fission-fusion dynamics and that the independence assumption for activity centers has been shown to be robust to departures (Reich and Gardner 2014).

We modeled the distribution of latent activity centers using an inhomogeneous point process (Borchers and Efford 2008) to estimate variation in cheetah density over the years. We expanded the 2-km resolution grid of the MMNR to include a 20-km buffer (Figure S1), which was large enough to ensure a negligible encounter probability at the edges (Royle et al. 2014); we also excluded the northwest escarpment, which was likely to have restricted cheetah movement (Broekhuis and Gopalaswamy 2016). The total state space, *S*, of the point process therefore included 1,381 discrete grid cells for a total area of 5,524 km^2^. The intensity of the point process (i.e., the expected density) within a grid cell *j* in a given year was a log-linear function:

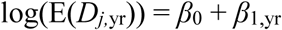

Here, *β*_0_ is the log-scale expected cheetah density in 2005, while *β*_1,yr_ is a vector of year-specific regression coefficients that estimate differences in expected density in later years (yr = 2013, 2014, 2015, or 2016). Conditional on the expected density for the year in which an individual was encountered (yr_*i*_), the probability of an individual’s activity center being located within a given grid cell was defined as:

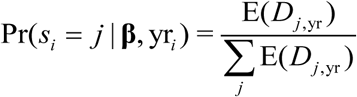

The marginal likelihood of the observations for each individual are then computed by integrating over all possible grid cells.

### Model fitting and sample period comparison

We fit the model using the multi-session sex-structured SCR framework in the R package oSCR (Sutherland et al. 2016) which maximizes the Poisson-integrated likelihood (Borchers and Efford 2008) and provides maximum likelihood estimates of model parameters. In addition to the parameters previously described, `oSCR` allows for estimating a sex ratio (ψ). Without specification of sex-specific parameters in the other SCR model components, estimates of ψ are derived entirely from the observed sex ratios of encountered individuals during each session (here, session = year).

We compared several sampling periods (early 5 months, full 10 months, late 5 months) to examine how differences in the observed data and parameter estimates affected population inferences. The early period spanned Jan–May and corresponded to a mostly hot and dry season that turns into long and heavy rains by May. The late period spanned Jun–Oct and corresponded to the cool season that follows the heavy rain season, during which widespread green vegetation supports a massive ungulate migration (Bell 1971, Sinclair and Norton-Griffiths 1995). This late period was similar to the 3-month sampling design (August–October) used by Broekhuis and Gopalaswamy (2016). The full 10-month sampling period spanned most of the year (Jan–Oct) and, while facilitating more observations and larger sample sizes of individuals and spatial encounters, was likely to violate the assumption of population closure to a greater degree than the 5-month periods. Aside from differences in the density estimates across time we were particularly interested in how other model parameters might change with variation in the number and type of spatial encounters, including the estimated NDVI relationships with encounter rate, sex ratios, and individual movement scale.

## Results

Monitoring efforts resulted in >7000 hours spent searching for and recording observations of cheetahs in the MMNR during 2005 and 2013–2016. The average number of hours searched each year was 1443 (range: 1086–1694) for the 10-month sampling period, which split into 623 (range: 513–790) for the early 5-month period and 820 (range: 573–989) for the late 5-month period (Table 1). Compared to either 5-month period, the increased sampling effort for the full 10 months always resulted in greater numbers (within a given year) of unique individuals encountered (median across years: full = 32, early = 20, late = 23), total encounter events (full = 101, early = 40, late = 60), and spatially distinct encounters (full = 58, early = 18, late = 28). The observed sex ratios were variable depending on the year and sampling period, though on a whole the median ratio was 1:1. We plotted the unique individuals encountered each year according to the midpoint ordinal date of their encounters, indicating the sampling period(s) in which they were observed (Figure 2). The patterns indicated similar ratios of females to males observed during all sampling period definitions.

**Table 1.** Summary of monitoring effort and adult cheetah encounters in the Masai Mara National Reserve during 2005 and 2013–2016. Results from the 3 sampling periods (early 5 months [Jan– May], full 10 months [Jan–Oct], and late 5 months [Jun–Oct]) include the hours spent searching, the number of unique individuals encountered (n) and broken down by sex (F/M), and the number of encounter events (y). Spatially distinct encounters occur across >1 grid cell and by definition involve recapture of an individual. For example, in 2005 there were 23 individuals encountered during the early 5-month period but only 8 (4 female; 4 male) were encountered in >1 grid cell.

| Months | Year | Hrs | Total encounters |  |  | Spatially distinct encounters |  |  |
| --- | --- | --- | --- | --- | --- | --- | --- | --- |
|  |  |  | n | (F/M) | y>0 | n | (F/M) | y>1 |
| 5 (Jan–May) | 2005 | 513 | 23 | (12/11) | 34 | 8 | (4/4) | 11 |
|  | 2013 | 790 | 12 | (6/6) | 34 | 6 | (3/3) | 18 |
|  | 2014 | 530 | 20 | (10/10) | 40 | 13 | (8/5) | 19 |
|  | 2015 | 705 | 24 | (11/13) | 68 | 12 | (6/6) | 39 |
|  | 2016 | 578 | 20 | (12/8) | 41 | 7 | (6/1) | 16 |
| 10 (Jan–Oct) | 2005 | 1086 | 26 | (14/12) | 47 | 14 | (7/7) | 21 |
|  | 2013 | 1535 | 20 | (10/10) | 73 | 13 | (9/4) | 46 |
|  | 2014 | 1465 | 34 | (14/20) | 112 | 26 | (12/14) | 73 |
|  | 2015 | 1694 | 32 | (13/19) | 142 | 19 | (10/9) | 92 |
|  | 2016 | 1438 | 32 | (17/15) | 101 | 22 | (12/10) | 58 |
| 5 (Jun–Oct) | 2005 | 573 | 11 | (7/4) | 13 | 2 | (0/2) | 2 |
|  | 2013 | 745 | 15 | (10/5) | 39 | 8 | (6/2) | 22 |
|  | 2014 | 935 | 30 | (13/17) | 72 | 17 | (5/12) | 39 |
|  | 2015 | 989 | 23 | (10/13) | 74 | 12 | (7/5) | 45 |
|  | 2016 | 860 | 27 | (14/13) | 60 | 14 | (9/5) | 28 |

**Figure 2.**
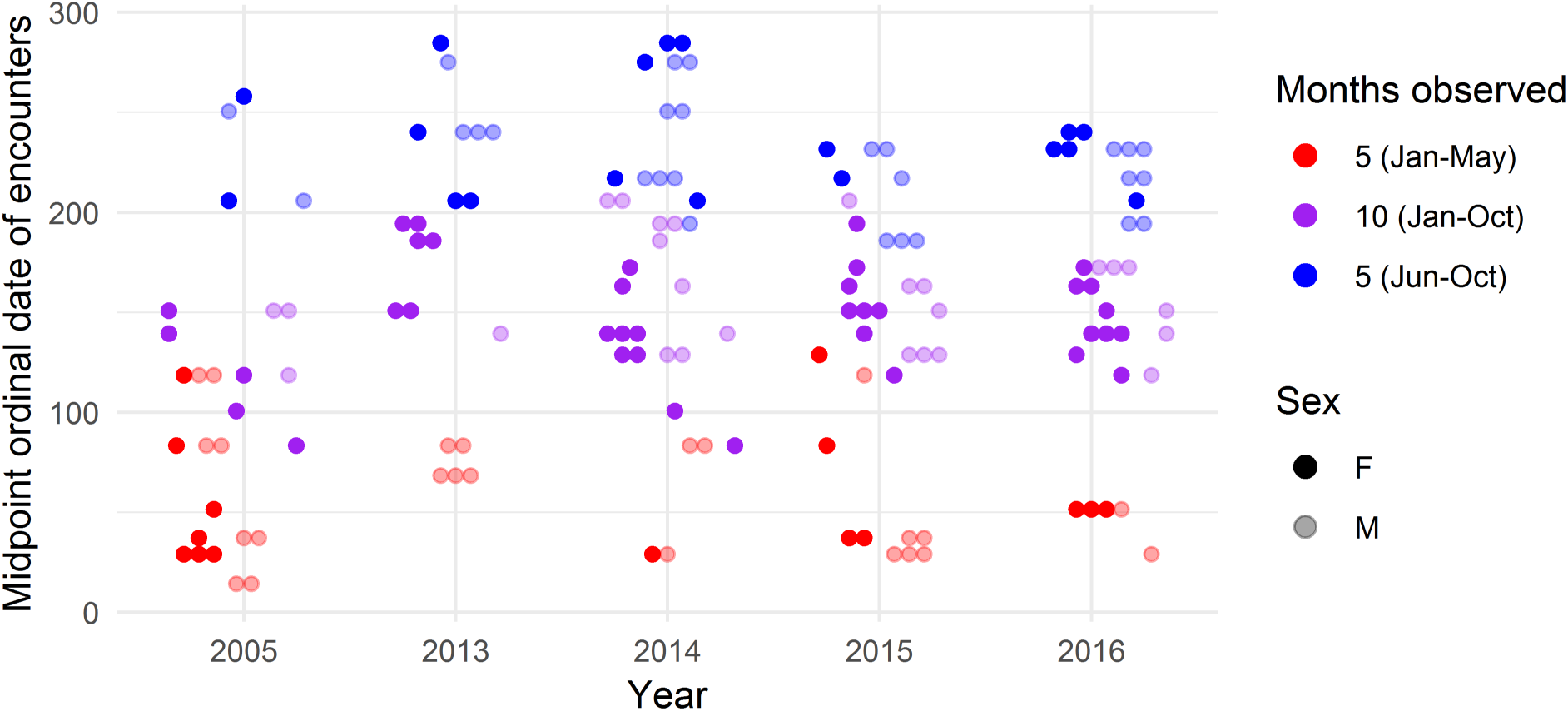
The midpoint ordinal date of encounter for each individual cheetah and the sampling periods in which they were encountered in the Masai Mara during 2005 and 2013–2016. Note, any individuals with encounters that spanned the full period (10 months) were included in the spatial capture-recapture models for all 3 periods.

The spatial capture-recapture models indicated similar patterns in density variation over time (Table 2–3; Figure 3), though fluctuations were mostly small relative to the uncertainty. The density estimates from 2005 had very large confidence intervals for the 5-month sampling periods due to small sample sizes. The full period density estimate (with 95% CI) for 2005 was 1.62 (1.02–2.57) cheetahs/100 km^2^. During 2013–2016, mean estimated density ranged from 0.60 (0.34–1.10) cheetahs/100 km^2^ in 2013 to 1.63 (0.97–2.73) cheetahs/100 km^2^ in 2014, and estimates matched closely across sampling periods within a given year. Precision of the density estimates was better for the 10-month sampling period, particularly with regards to the coefficients of variation (Table 3). Regardless of the sampling period, density estimates with a CV <0.30 could be achieved with >20 spatially distinct encounters (Figure 4).

**Table 2.** Parameter estimates from the spatial capture-recapture models of adult cheetah encounters in 2005 and 2013–2016 in the Masai Mara National Reserve, fit to data from the early period (Jan–May), full period (Jan–Oct), and late period (Jun–Oct). Estimates are on the scale of the appropriate link function, either log (***α***, ***β***) or logit (**ψ**).

| Process | $\theta$ | Effect | early (Jan–May) | | full (Jan–Oct) | | late (Jun–Oct) | |
| --- | --- | --- | --- | --- | --- | --- | --- | --- |
|  |  |  | Estimate | SE | Estimate | SE | Estimate | SE |
| Encounter | $\alpha_0$ | Intercept | −3.915 | 0.362 | −4.277 | 0.301 | −5.345 | 0.756 |
| | $\alpha_{1,2013}$ | Year | 1.261 | 0.459 | 0.784 | 0.319 | 1.732 | 0.790 |
| | $\alpha_{1,2014}$ | | 0.730 | 0.454 | 0.945 | 0.302 | 1.648 | 0.778 |
| | $\alpha_{1,2015}$ | | 1.089 | 0.408 | 0.998 | 0.303 | 1.660 | 0.782 |
| | $\alpha_{1,2016}$ | | 1.373 | 0.445 | 0.850 | 0.308 | 1.075 | 0.786 |
| | $\alpha_{2,2005}$ | NDVI×Year | 1.284 | 0.485 | 0.366 | 0.232 | −0.560 | 0.386 |
| | $\alpha_{2,2013}$ | | 1.167 | 0.393 | 0.709 | 0.252 | 0.621 | 0.344 |
| | $\alpha_{2,2014}$ | | −1.684 | 0.491 | −0.157 | 0.155 | 0.254 | 0.196 |
| | $\alpha_{2,2015}$ | | 0.961 | 0.335 | 0.775 | 0.219 | 0.580 | 0.297 |
| | $\alpha_{2,2016}$ | | −0.413 | 0.247 | 0.143 | 0.148 | 0.823 | 0.298 |
| | $\alpha_{3,2005}$ | NDVI<sup>2</sup>×Year | −0.920 | 0.397 | −0.369 | 0.225 | −0.245 | 0.382 |
| | $\alpha_{3,2013}$ | | −0.675 | 0.392 | −0.731 | 0.284 | −0.879 | 0.442 |
| | $\alpha_{3,2014}$ | | −1.569 | 0.458 | −0.609 | 0.174 | −0.636 | 0.230 |
| | $\alpha_{3,2015}$ | | −0.377 | 0.228 | −0.257 | 0.145 | −0.130 | 0.187 |
| | $\alpha_{3,2016}$ | | −0.408 | 0.219 | −0.127 | 0.125 | −0.236 | 0.211 |
| | $\alpha_4$ | Effort | 0.874 | 0.151 | 0.703 | 0.130 | 0.691 | 0.149 |
| | $\log(\sigma)$ | Dist. scale | 1.583 | 0.055 | 1.940 | 0.039 | 2.094 | 0.065 |
| Density | $\beta_0$ | Intercept | −2.243 | 0.291 | −2.732 | 0.236 | −2.760 | 0.682 |
| | $\beta_{1,2013}$ | Year | −1.474 | 0.418 | −0.668 | 0.328 | −0.791 | 0.735 |
| | $\beta_{1,2014}$ | | −0.491 | 0.389 | −0.146 | 0.294 | −0.147 | 0.709 |
| | $\beta_{1,2015}$ | | −0.867 | 0.358 | −0.415 | 0.295 | −0.659 | 0.714 |
| | $\beta_{1,2016}$ | | −0.904 | 0.379 | −0.316 | 0.297 | −0.246 | 0.714 |
| Sex ratio | $\psi_{2005}$ | Pr(M)×Year | −0.087 | 0.417 | −0.154 | 0.393 | −0.560 | 0.627 |
| | $\psi_{2013}$ | | 0.000 | 0.577 | 0.000 | 0.447 | −0.693 | 0.548 |
| | $\psi_{2014}$ | | 0.000 | 0.447 | 0.357 | 0.348 | 0.268 | 0.368 |
| | $\psi_{2015}$ | | 0.167 | 0.410 | 0.379 | 0.360 | 0.262 | 0.421 |
| | $\psi_{2016}$ | | −0.405 | 0.456 | −0.125 | 0.354 | −0.074 | 0.385 |

**Table 3.** Mean estimates (with standard errors and coefficients of variation) of cheetah density (#/100 km^2^) from the spatial capture-recapture models of adult cheetah encounters in 2005 and 2013–2016 in the Masai Mara National Reserve, fit to data from the early period (Jan–May), full period (Jan–Oct), and late period (Jun–Oct).

| Year | early (Jan–May) |  |  | full (Jan–Oct) |  |  | late (Jun–Oct) |  |  |
| --- | --- | --- | --- | --- | --- | --- | --- | --- | --- |
|  | Mean | SE | CV | Mean | SE | CV | Mean | SE | CV |
| 2005 | 2.65 | 0.77 | 0.29 | 1.63 | 0.38 | 0.24 | 1.58 | 1.08 | 0.68 |
| 2013 | 0.61 | 0.18 | 0.30 | 0.83 | 0.19 | 0.23 | 0.72 | 0.20 | 0.28 |
| 2014 | 1.62 | 0.43 | 0.26 | 1.41 | 0.25 | 0.18 | 1.37 | 0.28 | 0.20 |
| 2015 | 1.12 | 0.24 | 0.21 | 1.07 | 0.19 | 0.18 | 0.82 | 0.18 | 0.22 |
| 2016 | 1.07 | 0.27 | 0.25 | 1.19 | 0.22 | 0.18 | 1.24 | 0.27 | 0.22 |

**Figure 3.**
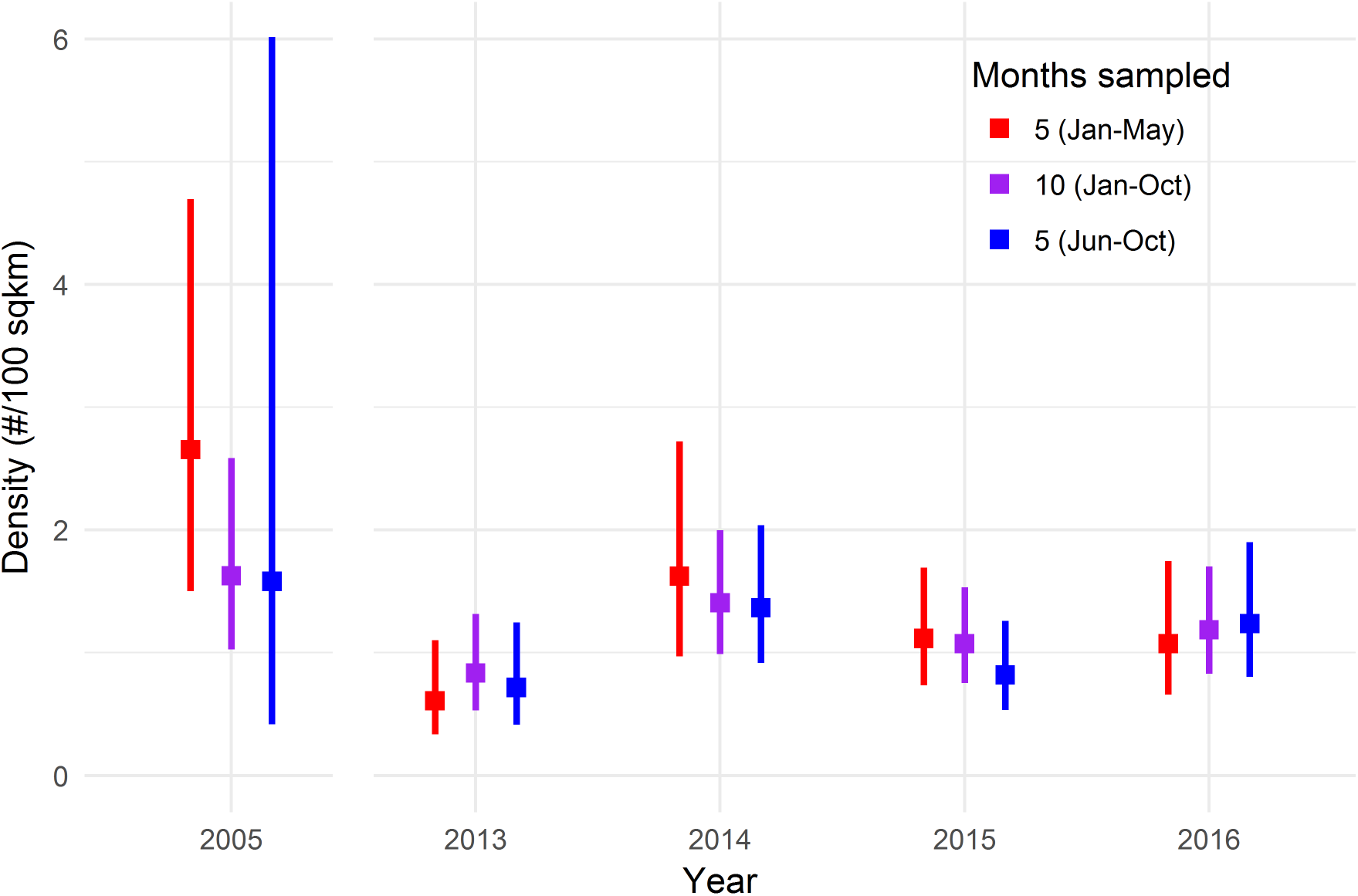
Mean estimates (with 95% CI) of cheetah density (#/100 km^2^) in the Masai Mara National Reserve in 2005 and 2013–2016 from spatial capture-recapture models fit using 5 months (early and late periods) and 10 months of surveys.

**Figure 4.**
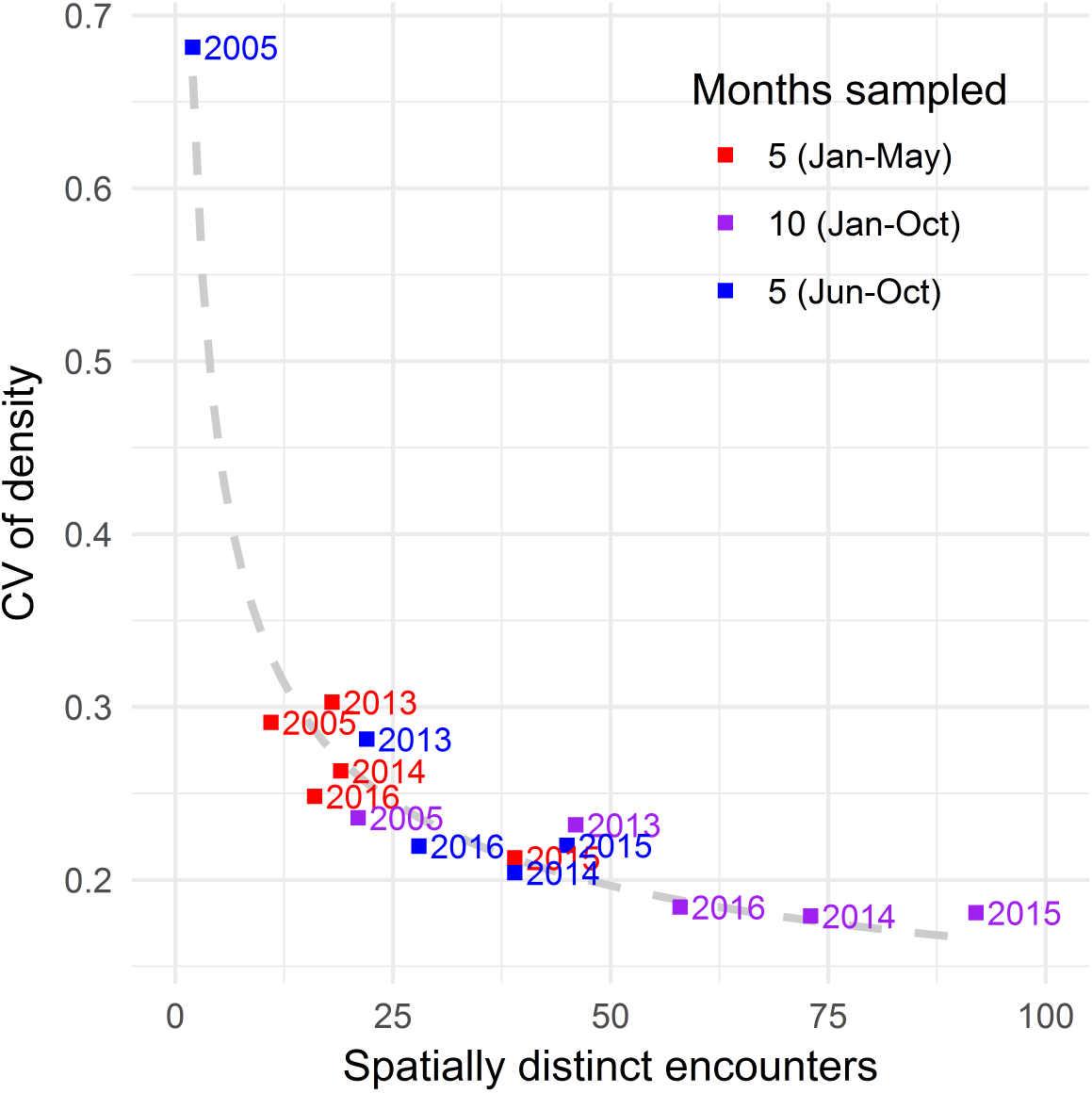
Relationship between number of spatially distinct encounters and coefficients of variation (CVs) for density estimates from the spatial capture-recapture models. Spatially distinct encounters occur across >1 grid cell and by definition involve recapture of an individual.

The relationships between encounter rate and NDVI variance were variable across years and across sampling periods within a year (Table 2; Figure 5). For most years and sampling periods, the maximum encounter rates occurred at mid to high values of relative NDVI variance. The early period in 2016 was the primary exception, suggesting higher encounter rates for cheetahs in low variance areas during Jan–May. The hours spent searching a grid cell (given the survey block within which it was located) had a strong positive relationship with encounter rate, and average encounter rates were higher in later years (2013–2016) compared to 2005. The scale parameter of the half-normal distance function was much smaller for the early 5-month period (*σ* = 4.9 km [4.4–5.4 km]) compared to the late 5-month period (*σ* = 8.1 km [7.2–9.2 km]) and the full 10-month period (*σ* = 7.0 km [6.4–7.5 km]).

**Figure 5.**
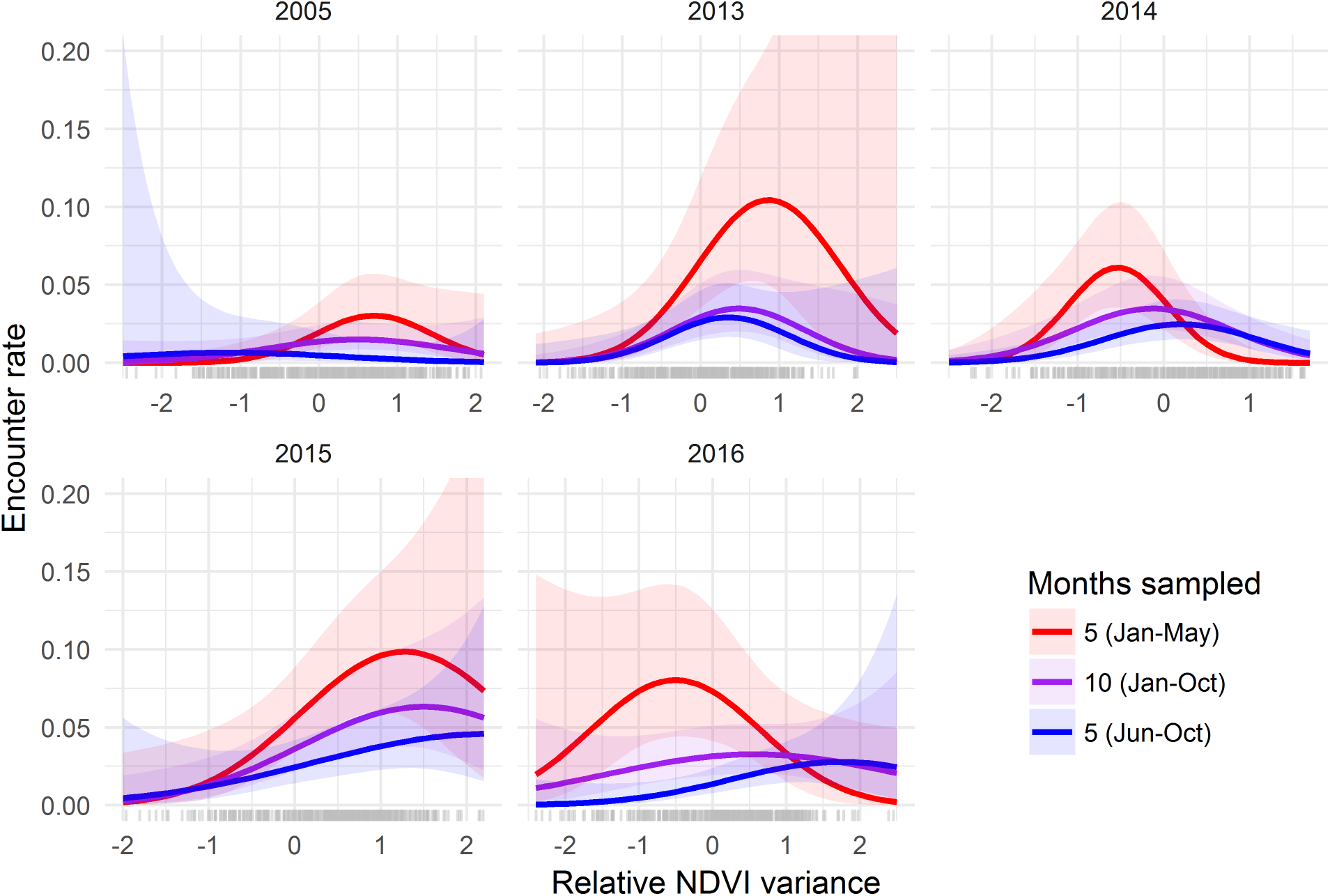
Predicted relationships (with 95% CI) between NDVI variance and cheetah encounter rate during 2005 and 2013–2016 from spatial capture-recapture models using 5 months (early and late periods) and 10 months of surveys. Values for NDVI variance were standardized to have mean 0 and unit variance within each year. Ticks at bottom indicate observed values at pixel locations within the MMNR.

The sex ratios were estimated to be largely even across all years and sampling periods as none of the logit-scale estimates were significantly different from 0 (Table 2), suggesting that the probability of an individual being a male did not vary considerably from 0.50.

## Discussion

Effective wildlife population monitoring spans enough time and space to detect change or variation that may require further investigation or be targeted for management action. In addition to adequate spatial and temporal extents, the sampling intensity needs to produce enough observations to ensure reasonable precision from statistical models designed to estimate population parameters. Our spatial capture-recapture models of cheetah encounters suggested little change in cheetah density from 2005 to 2013–2016 in the Masai Mara National Reserve, though there was some evidence that density fluctuated annually in recent years. The sampling period length (5 vs. 10 months) and timing (early, late, full year) over which spatial encounters were included in the modeling did not substantially alter inferences about density when sample sizes were adequate (e.g., ≥20 spatially distinct encounters). This suggests some flexibility in the design of search-encounter surveys for monitoring cheetahs over large landscapes.

We estimated an average cheetah density of ~1.2 cheetahs/100 km^2^, consistent with the impression that the MMNR provides important habitat for cheetahs in Africa. Cheetah density varies extensively throughout the current species range, from 0.02 cheetahs/100 km^2^ in areas of low productivity (Belbachir et al. 2015) to >2 cheetahs/100 km^2^ in the highly productive Serengeti (Durant et al. 2011, Durant et al. 2017). Broekhuis and Gopalaswamy (2016) used a similar search encounter design with SCR modeling and estimated a mean cheetah density of ~1.3 cheetahs/100 km^2^ in the MMNR and surrounding conservancies in 2014, which is consistent with our 2014 estimate from the late period (1.37 cheetahs/100 km^2^). Our additional years of monitoring indicated that density in some years may be nearly half that which was estimated in 2014.

Long-term studies of cheetah population trends in the Mara-Serengeti ecosystem have indicated a relatively stable density in recent years (Chauvenet et al. 2011, Durant et al. 2011). If the density fluctuation we estimated during 2013–2016 represents a real ecological phenomenon, as opposed to sampling variability, then our comparison with 2005 is difficult to interpret, given that this single year could have represented either ebb or flow for the cheetah population. Therefore, it is actually unclear whether cheetah density has declined in the MMNR during the past 10+ years. This uncertainty highlights the value of long-term monitoring programs, but also of monitoring designs that can estimate population size with useful precision. Our population modeling was limited to adult cheetahs and many individuals were encountered during only a portion of the year (Figure 2), therefore, population fluctuation in the MMNR is likely due to variable movement between the reserve and surrounding areas (e.g., Serengeti National Park). The magnitude of individual movements in cheetahs could make annual density an erratic statistic for an area the size of the MMNR (1510 km^2^), especially in the presence of non-resident, “floater” males (Caro 1994). Density estimation from SCR modeling is generally robust to transient individuals, though such movement dynamics could be explicitly modeled (Royle et al. 2016).

Based on the estimate of σ from the distance function, the mean 95% space use or home range area ranged from ~450 to ~1,200 km^2^ in the MMNR. Cheetah home ranges can be similar in size for males and females and overlap in areas where prey are non-migratory (Broomhall et al. 2003). In contrast, where ungulate prey are migratory, home ranges are comparatively larger with males forming small territories and females exhibiting roving behaviors (Caro 1994). Although there is a seasonal influx of migrant herbivores into the MMNR each year (Bell 1971, Stelfox et al. 1986, Sinclair and Norton-Griffiths 1995), resident herbivores are also present year-round in relatively high numbers. Thus, movements by cheetahs in the MMNR may be better predicted by interspecific competition with other large carnivores (Broekhuis et al. 2013) or the direct and indirect effects of people, rather than habitat suitability or prey populations. We caution any interpretation of the 95% space use approximation given the circular assumption of the bivariate normal distribution for *σ* (Royle et al. 2014). In addition, cheetah space use has been shown to be highly concentrated within a small portion of the home range (~14% of the total area), even for individuals that otherwise occupy large areas (Marker et al. 2008).

Several differences between our study and that of Broekhuis and Gopalaswamy (2016; hereafter, B&G) warrant discussion, given the similarity in our approaches to collecting and modeling spatial encounters of cheetahs in the Mara. First, B&G modeled the daily encounter probability over 90 days of sampling, while we summed our encounters over the relevant sampling period (5 or 10 months) and treated the counts as a Poisson random variable; given the low rates of encounter, these choices should have had a negligible influence (Royle et al. 2014). Second, our definitions of effort differed and B&G’s approach was preferable: using GPS tracks to define exactly which areas were searched. We did not have GPS track records for 2005 and instead attempted to systematically search pre-defined sections (i.e., blocks) of the MMNR for various lengths of time; such a definition of effort is approximate at best, though blocks were searched thoroughly when visited. Third, we observed a fairly even sex ratio of cheetahs that remained constant over the 5 years of surveys and is consistent with previous research in the Mara-Serengeti (Kelly et al. 1998). The extremely skewed ratio observed by B&G (F:M = 5:1) was potentially an artefact of a low sample size and short survey duration (3 months), though it should also be noted that most of their survey effort was in the conservancies to the north of the MMNR. Finally, B&G estimated a difference in the scale parameter (σ) between females and males; early data exploration here did not support such differences in our study, both given the observed maximum distances moved and preliminary estimates of σ from models with sex-specific parameters. Despite these differences, the close similarity in cheetah density estimates provides empirical support to the robustness of SCR modeling (Royle et al. 2014).

Improvements to the design of our search-encounter survey could make the effort more efficient and useful in other parts of the species range. We thinned almost 1/3 of our observed cheetah encounters before fitting the SCR models because of uneven effort across space and time. Ideally, areas would be searched with regular periodicity to ensure that inferences regarding individual movement matched in temporal scale at all spatial locations. This is typically the case for other common methods of collecting spatial encounters (e.g., camera trapping), where traps are operated on regular intervals (Royle et al. 2014). The problem of sampling regularity would be most acute for transient individuals; for example, 5 consecutive days of effort in a given location could yield a very different collection of encounters than 5 days spread across several months. Uneven spatial sampling makes the interpretation of posterior density surfaces from SCR models especially problematic and prone to artefacts (Efford 2018a), relegating the identification of “hot spots” (e.g., Broekhuis and Gopalaswamy 2016) to random error. Finally, the ability to traverse the landscape and get close enough to individuals for high quality photographs could limit the application of this survey to certain regions (e.g., protected areas). While long-range camera lenses may provide expanded opportunities for monitoring, it could still be difficult to clearly photograph both sides of every individual at great distances, ultimately increasing identification uncertainty (Augustine et al. 2018).

Other aspects of cheetah population ecology could be modeled with different or more complex analytical approaches to the individual encounter data we generated with the surveys. Our primary objective was a comparison between 2005 and 2013–2016, so we focused on understanding how best to estimate density within a given year (or seasonal period), while accommodating the sparse data from 2005. We hypothesized that individual space use and, thus, encounter probability would vary by habitat attributes and used NDVI variance as a proxy for ungulate habitat quality (Pettorelli et al. 2005, Bro-Jorgensen et al. 2008); in most years and seasonal periods, the spatial distribution of NDVI variance accounted for important variation in encounter rates. An open population model (Kendall et al. 1997) would allow for estimating survival and temporary emigration and potentially enable more comprehensive inferences than “snap-shot” density estimates (Harmsen et al. 2017). While open-population SCR models provide the opportunity to integrate spatial explicitness into estimation and prediction (e.g., Green et al. 2018b), the Bayesian frameworks typically used for fitting such models are notoriously slow and computationally demanding for complex spatiotemporal inferences. New approaches using maximum likelihood and hidden Markov models could provide promising alternatives (Glennie et al. 2017, Efford 2018b). Snap-shot estimates of population size across time are useful for wildlife monitoring, but understanding the mechanisms behind population changes can facilitate better conservation and management decision making (Harmsen et al. 2017).

## Acknowledgements

We thank the Kenya Wildlife Service, the Office of the President of Kenya, the Senior Warden of the Masai Mara National Reserve, and Mr. Brian Heath for the permissions to carry out this research. We also thank Kay Holekamp for providing assistance in the field. SMD was supported by grants from the Lakeside Foundation, Mr. Paul L. Davies II, and Dalbit Petroleum Ltd. DSG was supported by a Graduate Research Fellowship from the National Science Foundation. This research is presented in memory of Mr. Paul L. Davies II.

## Authorship

DWL, DSG, SMD, and EC designed the study. SMD, EC, and SM conducted the fieldwork, and DWL completed the modeling. All authors contributed to the manuscript writing.

## Data accessibility

Data will be archived with Dryad Digital Repository.

---

**Figure S1.**
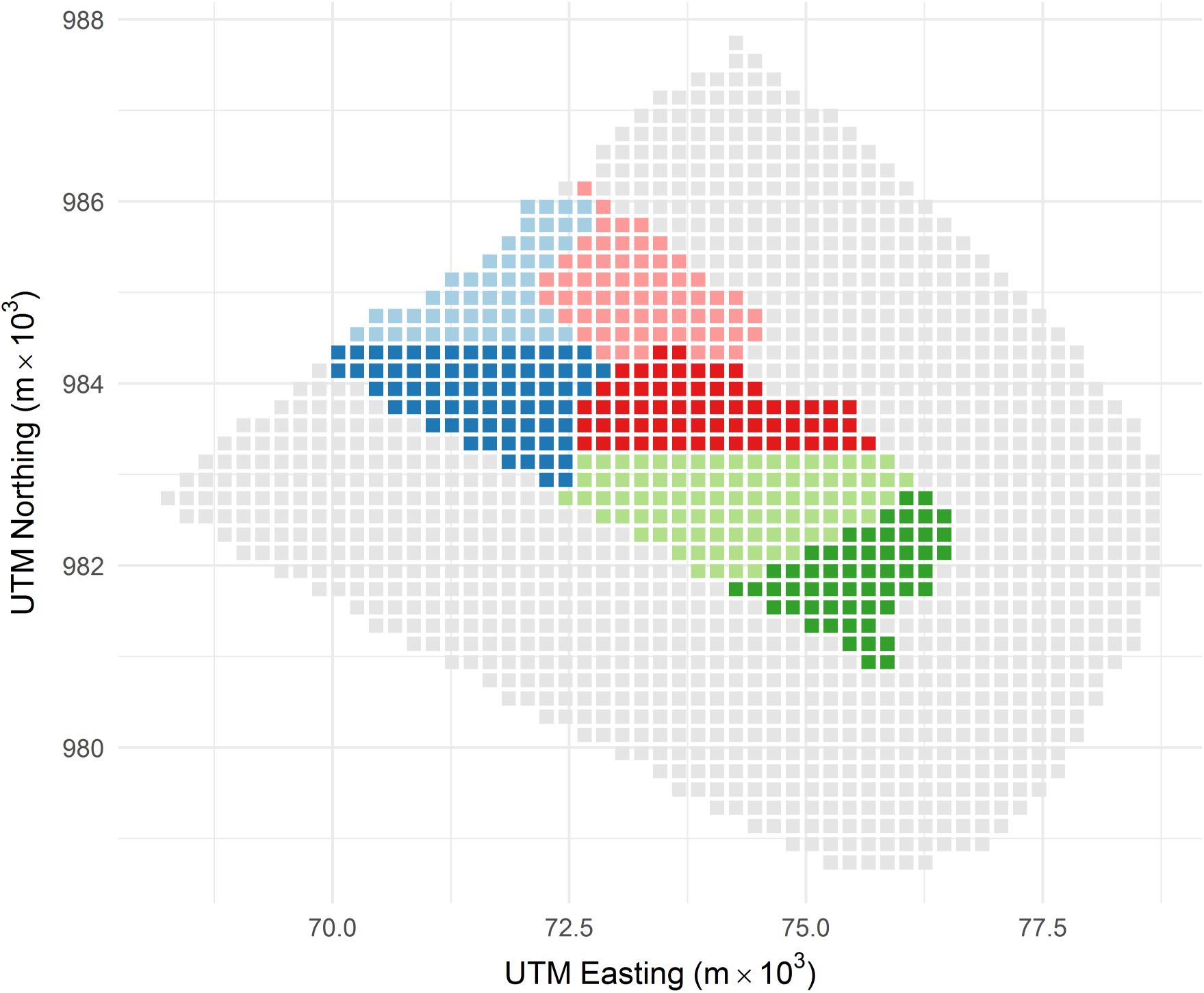
Grid cells illustrating state space used in the spatial capture-recapture models, with delineations of blocks according to survey effort. Light gray cells occurred in areas not searched but included as a buffer for population estimation.

**Figure S2.**
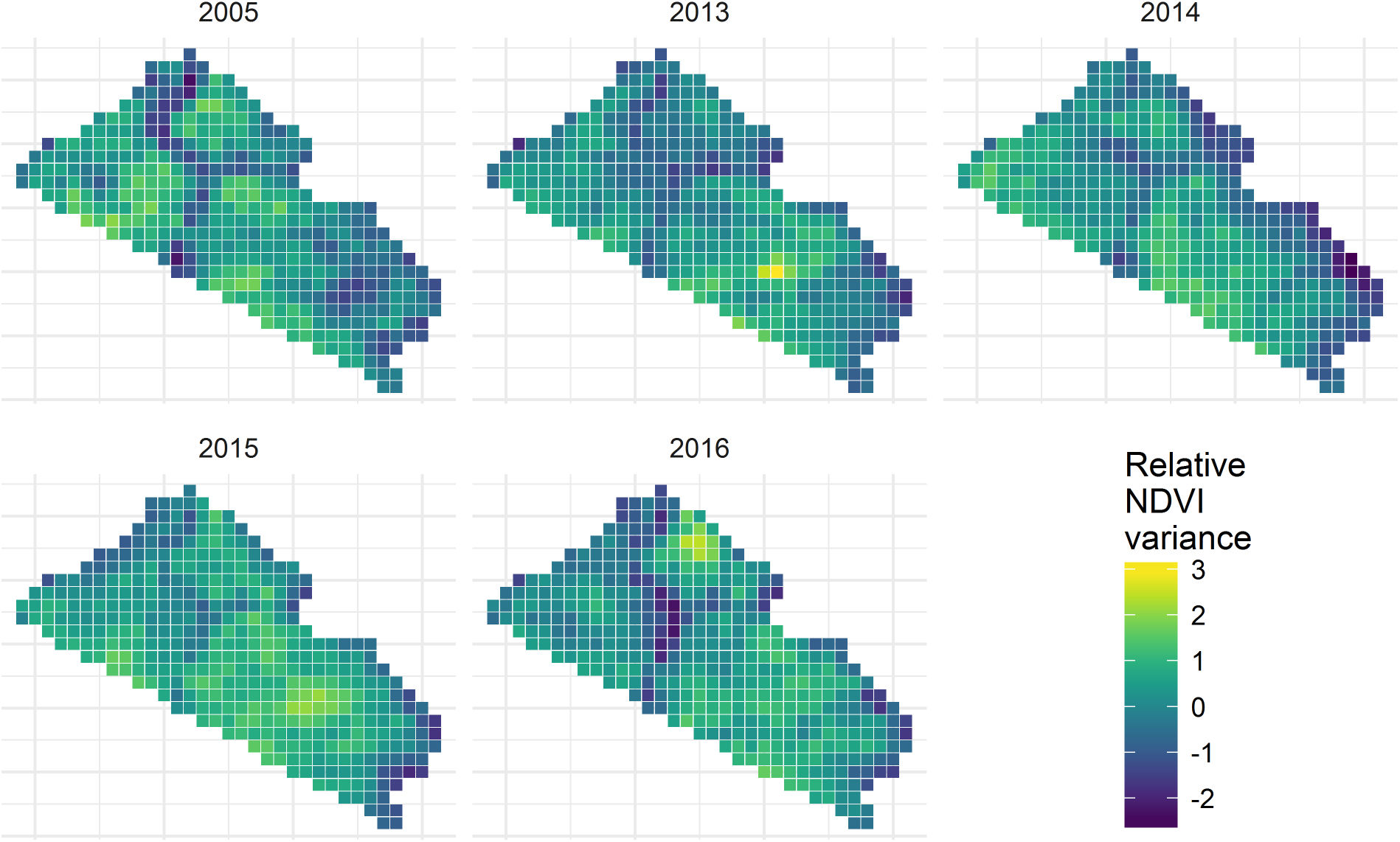
Spatial distributions of Normalized Vegetation Difference Index (NDVI) values in each year that served as covariates for encounter probability in the spatial capture-recapture models. Values represent the variance (standard deviation) in NDVI across the 36 satellite images (10-day intervals at 250 m resolution) for a given year. NDVI values were mean aggregated to the 2-km grid cells and standardized within each year to have mean of zero and unit variance. Satellite images acquired from the Famine Early Warning System Network hosted by the USGS/EROS Data Center (https://earlywarning.usgs.gov/fews/).

